# A novel multi SNP based method for the identification of subspecies and associated lineages and sub-lineages of the *Mycobacterium tuberculosis* complex by whole genome sequencing

**DOI:** 10.1101/213850

**Authors:** S Lipworth, R Jajou, A de Neeling, P Bradley, W van der Hoek, G Maphalala, M Bonnet, E Sanchez-Padilla, R Diel, S Niemann, Z Iqbal, G Smith, T Peto, D Crook, TM Walker, D van Soolingen

## Abstract

The clinical phenotype of zoonotic tuberculosis, its contribution to the global burden of disease and prevalence are poorly understood and probably underestimated. This is partly because currently available laboratory and *in silico* tools have not been calibrated to accurately identify all subspecies of the *Mycobacterium tuberculosis* complex (*Mtbc*). We here present the first such tool, SNPs to Identify TB (‘SNP-IT’). Applying SNP-IT to a collection of clinical genomes from a UK reference laboratory, we demonstrate an unexpectedly high number of *M. orygis* isolates. These are seen at a similar rate to *M. bovis* which attracts much health protection resource and yet *M. orygis* cases have not been previously described in the UK. From an international perspective it is possible that *M. orygis* is an underestimated zoonosis. As whole genome sequencing is increasingly integrated into the clinical setting, accurate subspecies identification with SNP-IT will allow the clinical phenotype, host range and transmission mechanisms of subspecies of the *Mtbc* to be studied in greater detail.

## Background

Tuberculosis (TB) is mostly caused by *Mycobacterium tuberculosis*, but also by other representatives of the *M. tuberculosis* complex (*Mtbc*). The *M. tuberculosis* complex is genetically highly conserved with the exception of *M. canetti*, almost exclusively restricted to patients from the Horn of Africa,^1^ which is genetically much more diverse. Indeed, it has been suggested *M. canetti* is the oldest divergent of the complex with the highest similarity to the common ancestor^2^.

The host range of the various subspecies differs; for instance *M. bovis* is more frequently observed in animals than humans, in whom it is traditionally thought of as a zoonosis secondary to unpasteurized dairy consumption^3^. Human infections are largely caused by the five main lineages or genotype families of *M. tuberculosis sensu stricto (Mtb)*, plus *M. africanum 1 and 2^4^*. ‘Animal’ strains include *M. bovis bovis, M. bovis* BCG, *M. bovis caprae, M. pinnipedii, M. suricattae, M. orygis, M. microti* and *M. mungi*, though several of these lack standing in the classical nomenclature ^5–10^.

The so-called ‘animal’ subspecies often grow poorly in culture, making drug-susceptibility testing difficult, and are not all well differentiated by commonly available PCR-based assays^11^. Their incidence and clinical significance is likely to be underestimated as a result. Routine whole-genome sequencing (WGS) of mycobacteria is now either underway or planned in an increasing number of settings^12^, providing the opportunity to better identify these subspecies and thereby improve our understanding of their clinical phenotype.

There is a notable geographic association with some subspecies of *Mtbc. M. canettii* for example is found almost exclusively in the Republic of Djibouti and the West African lineages (*Mtb* lineages 5 and 6) are strongly associated with West Africa^13^. Even within lineages there are some sub-lineages with a global distribution and others that are more restricted^14^. The reasons for this are not fully known but may include an interplay between host and/or pathogen genetics as well as historical geographical distribution in environmental/animal reservoirs.

From a clinical perspective there are conflicting reports regarding effects of *M. tuberculosis* lineage on clinical phenotype^15,16^. Genotypes vary in their association with multidrug and extensively drug resistant TB. A clear example of this is the Beijing genotype (later re-named lineage 2). In Asia, Europe and South Africa this sub-grouping has been significantly associated with transmission of resistant forms of TB^17^. In addition there is an apparent association between the lineage 2/Beijing strain and increased pathogenicity and virulence whilst *M. bovis subspecies bovis* and *M. canettii* are known to be naturally resistant to pyrazinamide^18,19^. It is therefore important to identify these as the causative organism at an early stage and adjust treatment accordingly. Other phenotypic differences between lineages include *in vitro* growth rates, hydrophobicity, clinical outcomes, transmissibility and host inflammatory response^20^.

In addition to differences in natural susceptibility to antituberculosis drugs, the identification of the subspecies of a *M. tuberculosis* complex isolate has important implications for contact investigations and source case finding. *M. bovis* BCG can cause complications usually, but not always, in immunosuppressed patients following BCG vaccination or be isolated as a result of bladder carcinoma treatment and should not be misdiagnosed as *M. tuberculosis*^21,22^*. M. microti* is found in wild cats and rodents and causes human infection which is usually in association with rodent contact^23^. *M. pinnipedii* is a bacterium causing tuberculosis in seals which is sometimes transmitted to humans during outbreaks in zoos or wildlife parks^24^. *M. orygis* is mostly isolated from gazelle species but in the past few years has also been seen in humans although how they contract this bacterium is still unclear^6^. The other animal subspecies are rarely encountered in wild animals. The specific clinical phenotype of animal strains is largely unknown at present. In order to facilitate studies to investigate this it is first necessary to develop a methodology to allow accurate identification.

Efforts to differentiate members of the *Mtbc* and study the phylogeny of the complex have thus far included analysis of large genomic deletions^25^, variable number tandem repeats (VNTR), spacer oligonucleotide typing, multilocus sequence analysis (MLST), and more recently single nucleotide based phylogenies^26^. Numerous tools now exist that make *in silico* predictions of lineages within the complex from WGS data, using a variety of approaches including the detection of single nucleotide polymorphisms (SNPs) from both unassembled and mapped genomes, comparison of de Brujin graphs, and minHash based comparisons^27–30^. None of these tools have yet been calibrated to reliably differentiate between all subspecies.

Whole genome sequencing has allowed greater understanding of the microbiological diversity within the complex. The diversity of clinical phenotypes associated with infection by animal lineages is largely unknown, partly because the identification of these organisms is currently difficult. Accurate identification of the causative subspecies in all cases would allow the burden of disease associated with animal lineages to be characterised and diversity in clinical phenotypes to be fully appreciated and subsequently better managed. A higher level of knowledge on the spread and host range of the subspecies would also provide a better basis on which to study the history of the evolutionary development of the complex as a whole. Here we use WGS data to identify SNPs that define each subspecies, lineage and sub-lineage within the *Mtbc* and use this SNP catalogue (implemented as ‘SNPs to Identify TB’, or ‘SNP-IT’) to characterize strains from a large collection of clinical isolates.

SNP-IT is freely available at github.com/samlipworth/SNP-IT

## Methods

### Calibration set

First, a set of isolates was defined from which to identify SNPs associated with subspecies, lineages and sub-lineages within the *Mtbc*. A training set of 323 isolates representing the five lineages of *Mtb*, plus *M. africanum 1&2*, was compiled from an archive of WGS at the University of Oxford. Sample selection was based on maximum likelihood (ML) phylogeny-assigned lineage (IQTREE-omp version 1.5.5 using a generalised time reversible model)^31^. In addition, all available animal strains from the collection at the National Institute for Public Health and the Environment (RIVM) in the Netherlands were added. This was supplemented with published strains for *M. suricattae, M. mungi* and the *dassie bacillus (ex Procavia capensis*) (supplement). These were identified using a combination of spoligotyping, RFLP, HAIN Genotype MTBC, and VNTR in accordance with current and previous standard practice.

### Bioinformatics

Parallel bioinformatics approaches were taken to assess applicability across pipelines. As such, reads from Illumina platforms were independently mapped to two different versions of the H37Rv reference genome. Reads were mapped to NC000962.3 with Breseq v0.28.1, using a minimum allele frequency of 80% and minimum coverage of 5, for SNP calls. Separately, reads were mapped again to NC000962.2, for which Snippy v3.1 was used with default settings (minimum coverage 10, minimum allele frequency 90%)^32,33^. We extracted all SNPs shared exclusively by members of each subspecies, lineage and sub-lineage. As the lineage-defining bases for lineage 4 as a whole do not vary from the reference genome, as it is itself lineage 4, we identified these positions by mapping a core SNP alignment to a maximum likelihood tree using Mesquite version 3.30^34^ These nucleotide loci were added to the catalogue of phylogeny-determining SNPs.

All newly sequenced genomes are available on NCBI under project accession number PRJNA418900.

### Validation set

To validate the algorithm, an independent collection of genomes (N = 516) was compiled using clinical isolates sequenced by Public Health England (PHE) Birmingham, identified as *Mtbc*, and not included in the calibration set. These were augmented with data from the public nucleotide archives to increase the representation of ‘animal’ subspecies (N = 102). To maximise inclusion of animal strains from the public archives, especially since we expected their identification to be problematic, we used the new Coloured Bloom Graph (CBG) software^35^. Using CBG, we searched a snapshot of the Sequence Read Archive (to Dec 2016, N = 455, 632) using our new set of reference kmers for Mykrobe predictor (see below). If CBG and SNP-IT agreed on the subspecies identity of the isolate, we included it in the validation set.

FASTA files (i.e. of the whole genome) were compared to the catalogue of phylogeny-determining SNPs to make predictions for PHE isolates, whereas for isolates downloaded from the nucleotide archive, only limited variant calling format (vcf) files were created using Snippy (i.e. only SNPs with respect to the reference genome were included to increase computational efficiency). To ensure genomic loci defining lineage 4 were included, a mutated reference genome was used to create these limited vcf files. SNPs in the query sample were compared to reference libraries of lineage specific SNPs for each clade. Query genomes were assigned to particular subspecies or lineage if at least 10% of lineage or subspecies specific SNPs were detected in the strain in question. All predictions were assessed against the ML phylogeny. For *M. mungi*, only one genome could be located in the public sequence libraries and so we could not validate this subspecies.

### Clinical isolates

To assess the caseload across the different members of the *Mtbc* seen by the Public Health laboratory in Birmingham, UK, we applied the algorithm to 3,128 *Mtbc* genome sequences from consecutively obtained clinical isolates. H37Rv is routinely sequenced by the laboratory on WGS plates; these isolates were not removed and their identification served as an internal control.

### Comparison to existing tools

Results were compared to those from existing software tools (KvarQ, Mykrobe Predictor and TB-Profiler). In order to allow our new data to be integrated with published SNP libraries^36^ and for practical reasons when modifying existing tools, we created a minimal SNP dataset. We filtered our larger SNP catalogue for synonymous SNPs which occurred in coding regions (as annotated by SnpEff version 4.3^37^) and selected one representative SNP for each subspecies, lineage and sub-lineage at random. The existing software packages were then modified to include reference SNPs (or kmers for Mykrobe predictor) for those subspecies, lineages or sub-lineages which they initially failed to identify.

## Results

### Calibration

In total, 13,544 SNPs were identified as being predictive of taxonomic and phylogenetic groups of interest (median of 265 SNPs per group, inter quartile range 345).

### Validation

All predictions made by SNP-IT across all of the subspecies, lineages and sub-lineages were consistent with the ML phylogeny (Table 1).

**Table 1:**
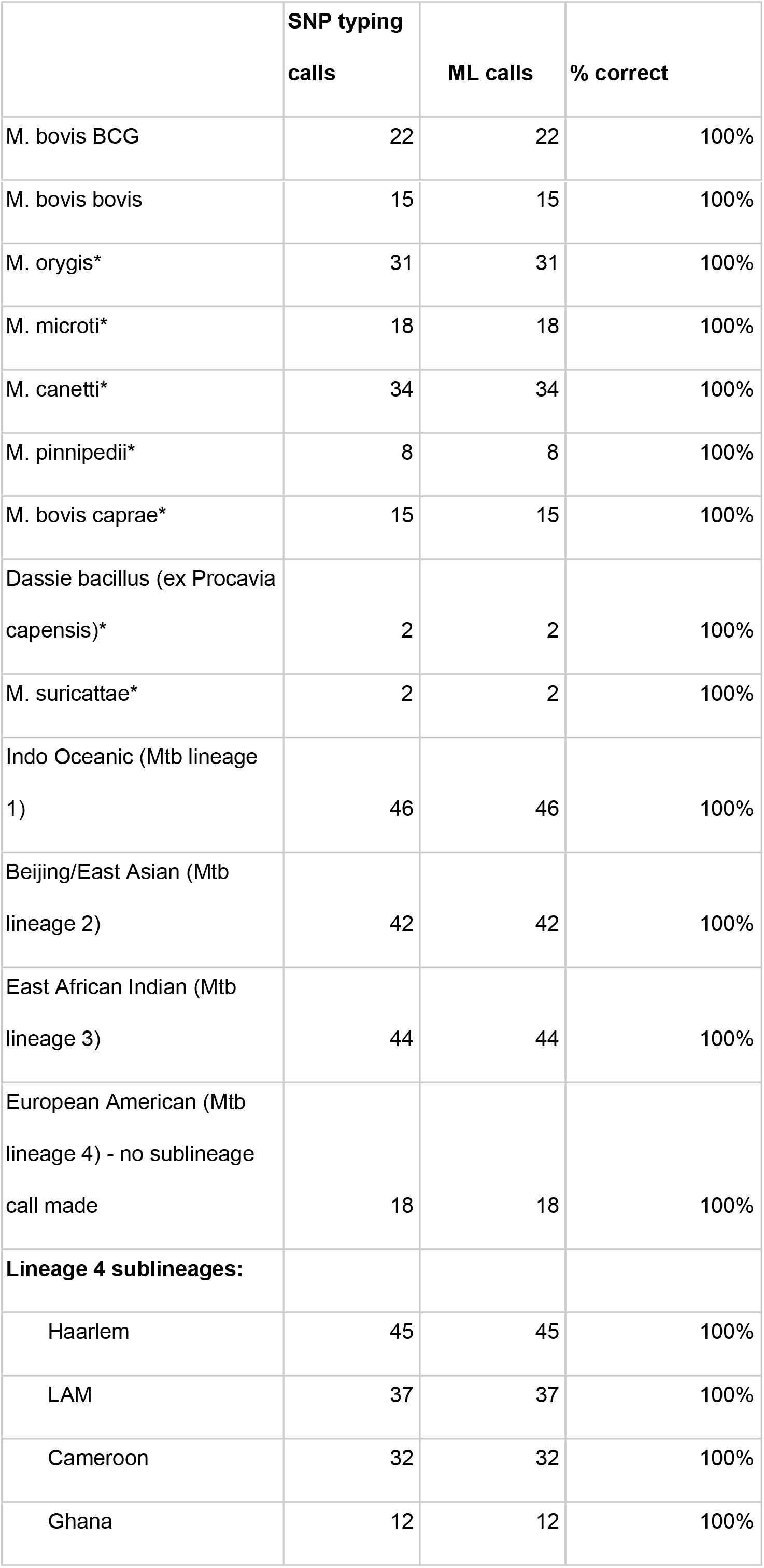
comparison between speciation calls for the validation set made by SNP-IT and position on maximum likelihood phylogenetic tree. * Validation set for subspecies augmented with published strains.

|  | SNP typing<br>calls | ML calls | % correct |
| --- | --- | --- | --- |
| M. bovis BCG | 22 | 22 | 100% |
| M. bovis bovis | 15 | 15 | 100% |
| M. orygis* | 31 | 31 | 100% |
| M. microti* | 18 | 18 | 100% |
| M. canetti* | 34 | 34 | 100% |
| M. pinnipedii* | 8 | 8 | 100% |
| M. bovis caprae* | 15 | 15 | 100% |
| Dassie bacillus (ex Procavia capensis)* | 2 | 2 | 100% |
| M. suricattae* | 2 | 2 | 100% |
| Indo Oceanic (Mtb lineage 1) | 46 | 46 | 100% |
| Beijing/East Asian (Mtb lineage 2) | 42 | 42 | 100% |
| East African Indian (Mtb lineage 3) | 44 | 44 | 100% |
| European American (Mtb lineage 4) - no sublineage call made | 18 | 18 | 100% |
| **Lineage 4 sublineages:** |  |  |  |
| Haarlem | 45 | 45 | 100% |
| LAM | 37 | 37 | 100% |
| Cameroon | 32 | 32 | 100% |
| Ghana | 12 | 12 | 100% |
| S-type | 45 | 45 | 100% |
| Tur | 25 | 25 | 100% |
| Uganda | 22 | 22 | 100% |
| Ural | 19 | 19 | 100% |
| X-type | 45 | 45 | 100% |
| West African 1 (Mtb lineage 5) | 13 | 13 | 100% |
| West African 2 (Mtb lineage 6) | 11 | 11 | 100% |
| Ethiopian (Mtb lineage 7) | 15 | 15 | 100% |

Within the validation set we included 18 genomes from lineage 4 for which no further sub-lineage could be assigned by SNP-IT and which we therefore defined by the presence of conserved bases relative to the reference. Their position on a ML tree revealed that these were phylogenetically distinct from other currently named sub-lineages of lineage 4. We therefore carried out a *post hoc* analysis of all genomes in the validation and clinical sets labelled by SNP-IT as being ‘lineage 4’ with no further sub-lineage name given, comparing their position on the ML tree to the nomenclature of Coll^36^ and Stucki^38^ (supplementary figure 1). Of 886 genomes labelled as ‘lineage 4’ with no further sublineage identification by SNP-IT, the vast majority were lineage 4.10 (817) (Stucki) also known as lineage 4.7 (35)/4.8 (247)/4.9 (535) (Coll). A minority belonged to lineage 4.5 (35) and lineage 4.6 (27) where they were phylogenetically distinct from the Uganda and Cameroon sub-lineages. The remainder (7) were not further classifiable from lineage 4 by Coll nomenclature. We subsequently updated our SNP catalogue for lineage 4 to include these unnamed sub-lineages using the same methods as we described for the original training set. 368 samples (all belonging to lineage 4.9) were found to be H37Rv when we unblinded ourselves to their corresponding lab records.

### Phylogenetic SNPs in drug resistance associated genes

Using a previously published list of drug resistance associated genes for *M. tuberculosi*s^39^, we searched all animal subspecies for phylogenetic SNPs in drug associated genes (supplementary table 3). All animal subspecies contain phylogenetic SNPs in these genomic regions, but on the basis of our data we were unable to determine whether any of these mutations are linked to lineage specific resistance as we did not have the corresponding phenotypic drug susceptibility testing data.

### Determining the prevalence of *Mtbc* cases in a collection of clinical isolates

We retrospectively applied SNP-IT to clinical isolates sequenced as part of the routine PHE diagnostic workflow in Birmingham to estimate the prevalence of the animal subspecies amongst *Mtbc* samples. Of 3,128 samples for which there was a whole-genome sequence available, 24 were identified as *M. orygis*, three as *M. microti*, 34 as *M. bovis*, and one as *M. caprae*. We did not identify any *M. pinnipedii, M. canettii* nor *Mtb* lineage 7 (*Aethiops vetus;* Ethiopian) isolates. Full results are depicted in figure 3/table 2 below.

**Figure 1:**
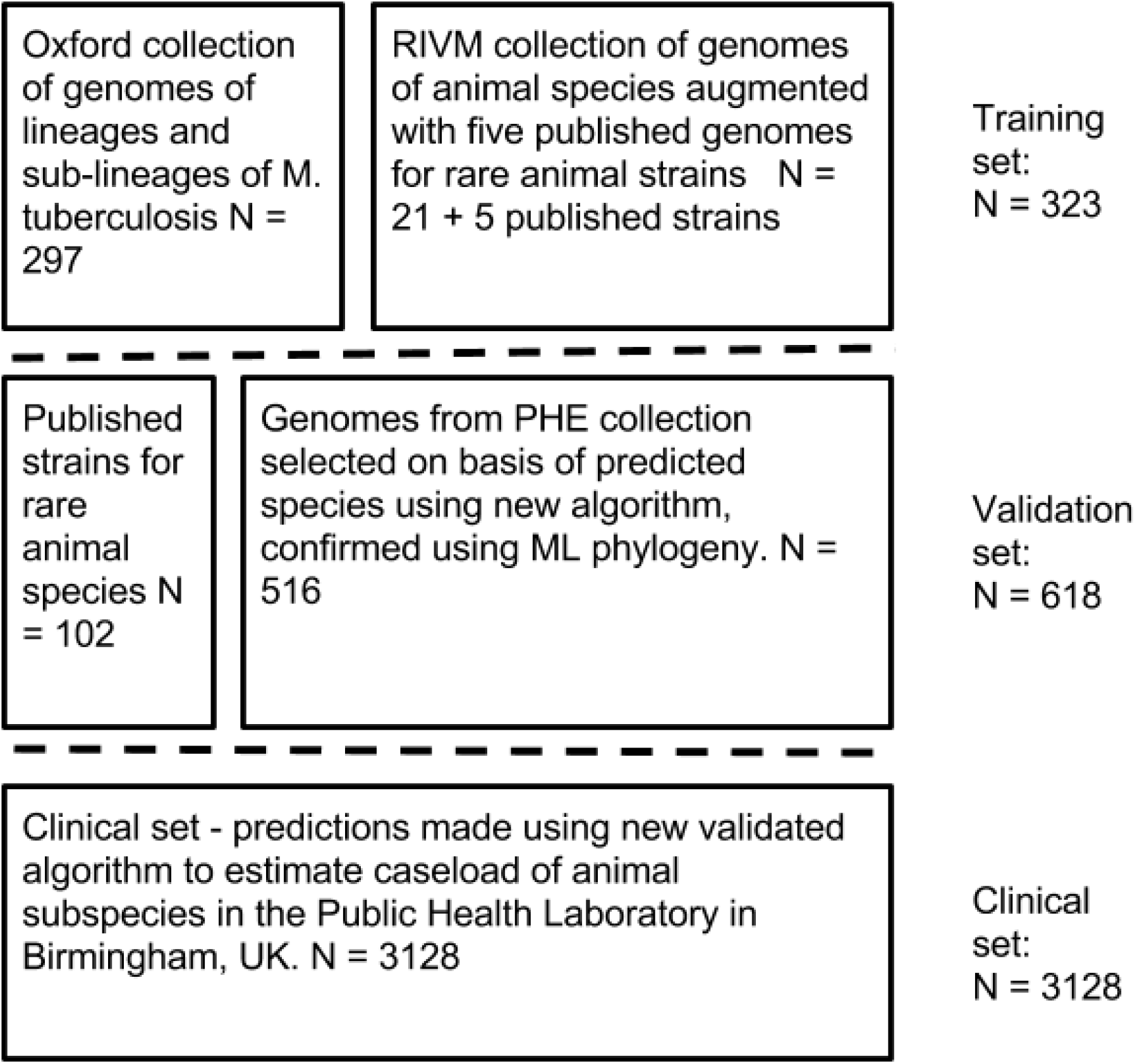
Description of the datasets used in the three stages of training, validation and application of the new algorithm.

**Figure 2:**
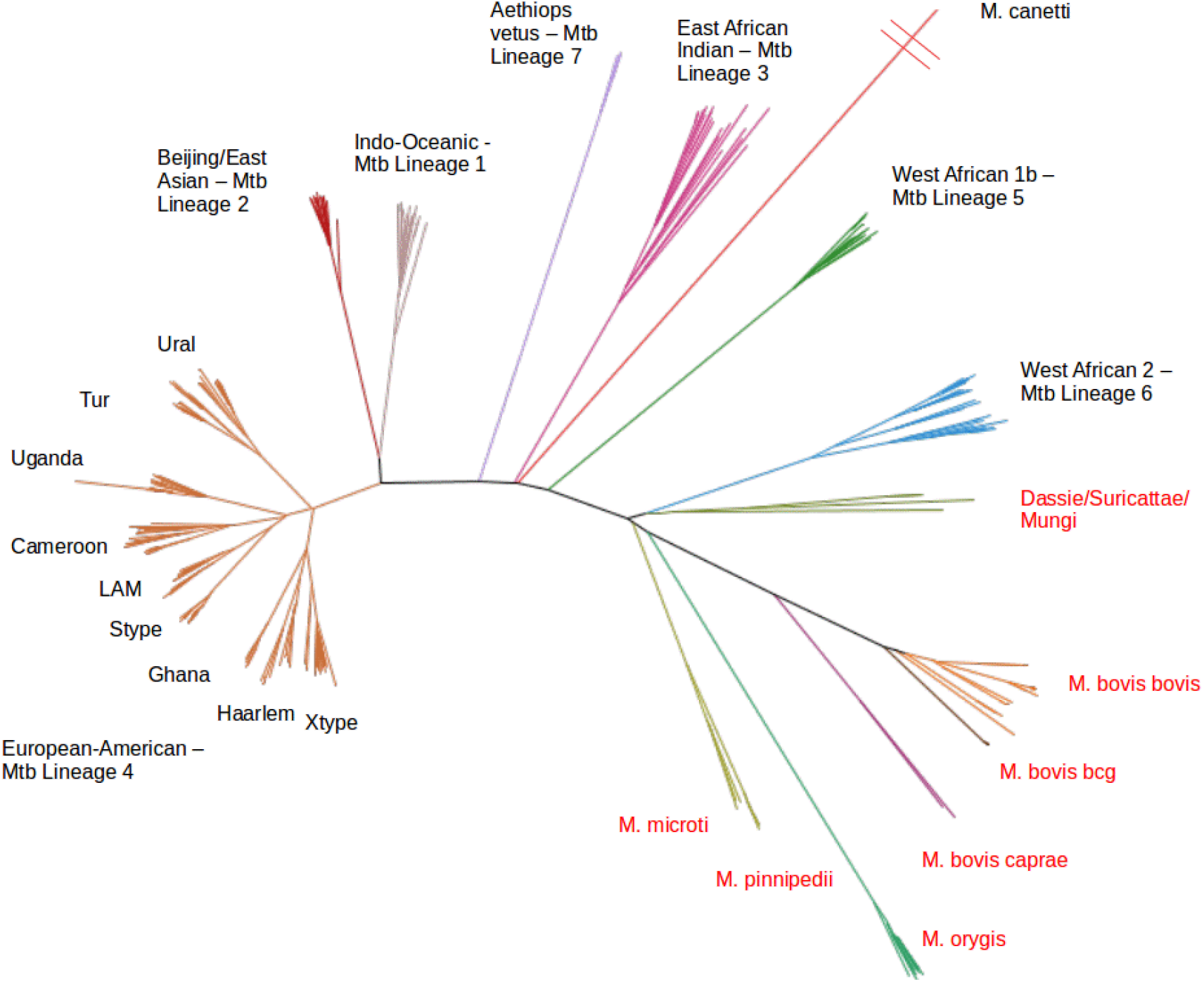
Maximum likelihood tree of all 323 samples in the training set. Red coloured text denotes animal subspecies.

**Figure 3:**
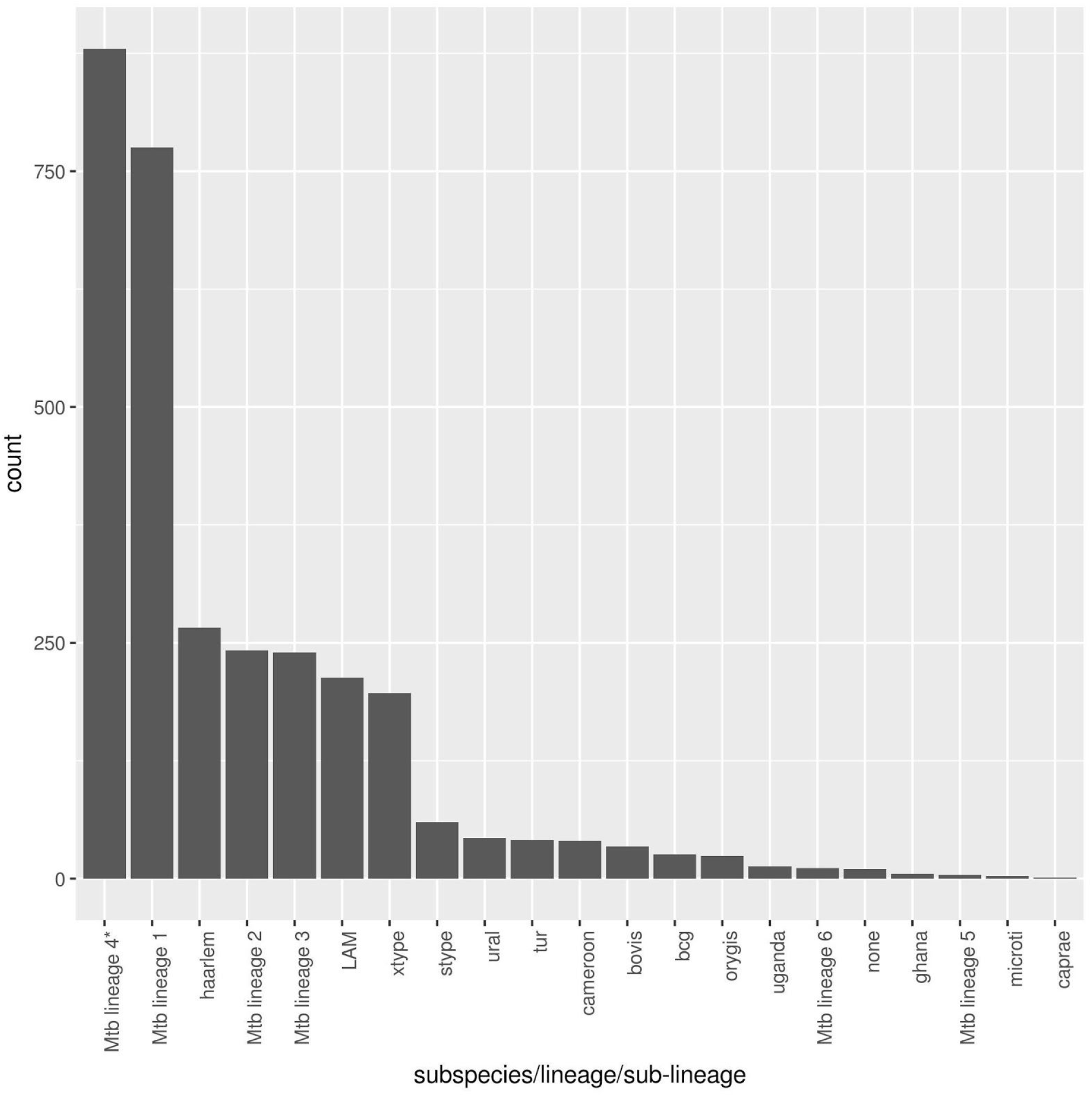
count of all lineages, sub-lineages and subspecies in the Birmingham clinical isolate collection as determined by our new typing dataset. ‘Lineage 4’ describes isolates that were not identified as belonging to one of the named sub-lineages, 368/880 were H37Rv which is commonly re-sequenced by the lab.

**Table 2:** Speciation predictions for the collection of clinical isolates using our new SNP-IT approach. * these samples belong to lineage 4 but not to one of the named sub-lineages, 368/880 were H37Rv which is commonly re-sequenced by the lab.

| **Subspecies/Lineages** | **N.** |
| --- | --- |
| *M. bovis* (BCG) | 26 |
| *M. bovis* | 34 |
| Indo Oceanic (Mtb lineage 1) | 775 |
| Beijing/East Asian (Mtb lineage 2) | 242 |
| East African Indian (Mtb lineage 3) | 240 |
| European American (Mtb lineage 4) - no | 880 |
| sub-lineage call made* |  |
| X-type | 197 |
| LAM | 213 |
| Ghana | 5 |
| Uganda | 13 |
| Ural | 43 |
| S-type | 60 |
| Tur | 41 |
| Haarlem | 266 |
| Cameroon | 40 |
| West African 1 (Mtb lineage 5) | 4 |
| West African 2 (Mtb lineage 6) | 11 |
| No call made | 10 |
| *M. orygis* | 24 |
| *M. microti* | 3 |
| *M. bovis caprae* | 1 |
| **Total** | **3128** |

### Updating existing software

A recent review found that KvarQ and TB-Profiler had a 100% accuracy for lineage prediction, compared to 93.4% for Mykrobe predictor^40^. Given each of these systems uses the same principle of lineage defining SNPs to predict lineage, the differential performance is likely to stem from the SNP catalogues deployed. We therefore modified the SNP catalogues of KvarQ. Mykrobe predictor and TB-Profiler using our minimum (single) SNP catalogue and re-ran the software on our collection of clinical isolates.

Strain characterisation by our new ‘SNP-IT’ typing system was first compared to that of KvarQ, TB-Profiler and Mykrobe predictor on default settings. Compared to SNP-IT, Mykrobe predictor made no mistakes assigning strains to lineages 1 to 4. The lineage 4 associated sub-lineages however could not be identified by Mykrobe predictor. Additionally, it did not differentiate lineage 5 from lineage 6, and all *M. orygis* samples were solely reported as West African. *M. pinnipedii* was reported as *M. microti*. KvarQ identified lineages 1 to 6 correctly but reported everything else as ‘animal lineage’. TB-Profiler reported lineages 1 to 5 correctly but made a single miscall for a lineage 6 sample, which it reported as *M. bovis*. It was furthermore unable to delineate among ‘animal’ subspecies which were all reported as *M. bovis/M. africanum*.

After modifying the KvarQ, TB-Profiler and Mykrobe predictor databases to our minimal SNP catalogue, all systems identified all of the subspecies, lineages and sub-lineages with 100% accuracy. SNP-IT was unable to identify 10 samples because less than 10% of type-specific SNPs were present in these strains. This was because our pipeline made no call at the lineage informative sites due to the presence of a minor allele, most likely due to contamination or a mixture of two strains in the sample. TB-Profiler and Mykrobe predictor were however both able to identify two of these isolates as mixed Beijing strain/M. *orygis* using our new minimal SNP dataset.

## Discussion

SNP typing is a powerful method for discriminating between subspecies of the *M. tuberculosis* complex, which are often not discernible by conventional laboratory methods such as the Reverse Line Blot Methodology. The SNP databases of Coll, Stucki and Comas are currently used as the knowledge base for KvarQ, TB-Profiler and Mykrobe predictor^36,41,42^. None of these however provide adequate resolution for the animal lineages. In contrast, SNP-IT is able to assign lineage to all samples tested with 100% accuracy. Implementing this fine-resolution algorithm into a routine diagnostic work flow would be a major step towards understanding the epidemiology and pathogenicity of the less common members of the *Mtbc*. It would furthermore provide an important basis for studies on the development and the phylogeny of the complex. The global burden of zoonotic tuberculosis, largely caused by *Mycobacterium bovis*, is thought to be both underestimated and increasing^43^. In the UK approximately 1% of tuberculosis cases are caused by *M. bovis*, mostly due to re-activation in older individuals from previous exposure to unpasteurised milk products or travel/migration, although human to human transmission clusters have been identified^44^. This number is likely to be higher in endemic areas where humans have closer contact to animals, however accurate assessment of prevalence is made difficult by a lack of diagnostic tools and surveillance in these areas^45^. Even where routine diagnostic whole genome sequencing is available and *M. bovis* can be reliably distinguished from *Mtb*, there is at present an inability to discriminate between *M. africanum, M. bovis* and the other animal lineages using existing SNP typing systems.

By applying our system to a clinical dataset we discovered a higher than expected number of *M. orygis* samples. This recently described member of the complex has a host range which includes oryxes, waterbucks, gazelles, rhesus monkeys, cows and rhinocerae^6,46,47^. Several human cases have been described in Dutch patients of South/South East Asian origin^6^ but there have not been any cases described in the UK. Human to animal transmission has been described in a single case in New Zealand^48^.

In order to recognise the particulars of the clinical phenotype, epidemiology and optimal management strategy of *M. orygis* infection it is firstly important to accurately distinguish these cases from *M. africanum*. This is currently not possible by either the Hain Genotype MTBC probe or existing SNP based platforms. Given the large resource aimed at controlling bovine tuberculosis, it is interesting that another zoonosis is seen at a similar rate in this collection of clinical isolates. This raises questions about the host range of *M. orygis* in the UK, and how these individuals came to be infected: questions with potential implications for TB control both in animals and humans. From an international perspective it is likely that next to *M. bovis, M. orygis* plays an important and currently poorly understood role in zoonotic transmission in Africa and other high prevalence settings with extensive animal contact.

There were three clinical isolates of *M. microti*, all of which were also correctly identified by Mykrobe predictor but not KvarQ or TB-Profiler with their default reference libraries. There have been at least 26 cases of *M. microti* infection in humans reported in the literature both with and without close contact with relevant animals being a feature of the clinical history^23,49–53^. In a case series from Scotland, three out of four *M. microti* infections occurred in individuals not known to be immunocompromised and two of these had no known contacts with relevant animals^51^. In contrast to *M. bovis*, most infections have been reported to cause pulmonary tuberculosis which raises the possibility of onward transmission though this has not yet been reported.

In order to demonstrate that there is no material difference between existing platforms other than the SNP databases which they use, we adjusted their reference databases and re-ran the programmes on a collection of clinical isolates. We here demonstrated that all three systems tested (KvarQ, Mykrobe predictor and TB-Profiler) are identical in performance when given the same SNP reference database. There are differences between the packages including the user interface, method applied (eg mapped vs. unmapped calling), programming architecture, ability to call mixed samples and speed which ultimately come down to user preference and end application. We demonstrate here however that the clinically meaningful differences highlighted in a recent review are easily ameliorated by improving the underlying dataset^40^.

In all of the ‘animal’ lineages there are phylogenetic SNPs in drug resistance associated genes. Where these are not associated with drug resistance, they can be helpfully annotated as such by diagnostic algorithms and set aside for purposes of predicting susceptibility. An unavoidable weakness of any SNP-based approach is its vulnerability to null-calls due to minor alleles at lineage-informative positions, or a lack of coverage. SNP-IT uses the entire library of lineage defining SNPs such that this is not an issue unless it occurs at the majority of lineage informative positions.

In conclusion, in this study we present SNP-IT, which is the first SNP typing system to identify all subspecies, lineages and sub-lineages with 100% accuracy within the *Mtbc*. When the knowledge base behind SNP-IT is applied to three existing software packages, each performs equally well. By demonstrating the practical application of this knowledge base to a routinely collected clinical dataset, we reveal an unexpectedly high prevalence of *M. orygis*, providing an opportunity to further explore the clinical phenotype of this infection, and that of other animal lineages, in the future. As more healthcare systems begin to routinely use whole genome sequencing, there is an opportunity to accurately diagnose the causative subspecies of TB in all cases. This will allow previously unrecognised zoonosis and reverse zoonosis to be identified and control interventions to be implemented in the interests of ‘One Health’.

## Author Contributions

Contributed to drafting of manuscript: SIWL, TMW, DvS, RJ

Bioinformatic analysis and programming: SIWL, RJ, AdN, ZI, PB

Study design and analysis plan: TMW, RJ, SIWL, TP, DC, AdN, DvS

Sample collection and data acquisition: TMW, DvS, RJ, GS, GM, MB, ESP, RD, SN

All authors reviewed the draft manuscript and approved the final version for publication

## Materials and correspondance

Dr Sam Lipworth

## References

1. van Soolingen, D. et al. A novel pathogenic taxon of the Mycobacterium tuberculosis complex, Canetti: characterization of an exceptional isolate from Africa. Int. J. Syst. Bacteriol. 47, 1236–1245 (1997).

2. Fabre, M. et al. High genetic diversity revealed by variable-number tandem repeat genotyping and analysis of hsp65 gene polymorphism in a large collection of ‘Mycobacterium canettii’ strains indicates that the M. tuberculosis complex is a recently emerged clone of ‘M. canettii’. J. Clin. Microbiol. 42, 3248–3255 (2004).

3. Grange, J. M. Mycobacterium bovis infection in human beings. Tuberculosis 81, 71–77 (2001).

4. Comas, I. et al. Out-of-Africa migration and Neolithic coexpansion of Mycobacterium tuberculosis with modern humans. Nat. Genet. 45, 1176–1182 (2013).

5. Sakai, S. & Imachi, H. Methanocellales. in Bergey’s Manual of Systematics of Archaea and Bacteria (John Wiley & Sons, Ltd, 2015).

6. van Ingen, J. et al. Characterization of Mycobacterium orygis as M. *tuberculosis complex subspecies*. Emerg. Infect. Dis. 18, 653–655 (2012).

7. Parsons, S. D. C., Drewe, J. A., Gey van Pittius, N. C., Warren, R. M. & van Helden, P. D. Novel cause of tuberculosis in meerkats, South Africa. Emerg. Infect. Dis. 19, 2004–2007 (2013).

8. Mostowy, S., Cousins, D. & Behr, M. A. Genomic interrogation of the dassie bacillus reveals it as a unique RD1 mutant within the Mycobacterium tuberculosis complex. J. Bacteriol. 186, 104–109 (2004).

9. Alexander, K. A. et al. Novel Mycobacterium tuberculosis complex pathogen, M. mungi. Emerg. Infect. Dis. 16, 1296 (2010).

10. Fabre, M. et al. Molecular characteristics of ‘Mycobacterium canettii’ the smooth Mycobacterium tuberculosis bacilli. Infect. Genet. Evol. 10, 1165–1173 (2010).

11. Hain Lifescience. GenoType MTBC | Differentiation of the Mycobactrium tuberculosis complex from culture Available at: http://www.hain-lifescience.de/en/products/microbiology/mycobacteria/tuberculosis/genotype-mtbc.html. (Accessed: 19th September 2017)

12. Walker, T. M. et al. Tuberculosis is changing. Lancet Infect. Dis. 17, 359–361 (2017).

13. Gagneux, S. & Small, P. M. Global phylogeography of Mycobacterium tuberculosis and implications for tuberculosis product development. Lancet Infect. Dis. 7, 328–337 (2007).

14. Koeck, J.-L. et al. Clinical characteristics of the smooth tubercle bacilli ‘Mycobacterium canettii’ infection suggest the existence of an environmental reservoir. Clin. Microbiol. Infect. 17, 1013–1019 (2011).

15. Pareek, M. et al. Ethnicity and mycobacterial lineage as determinants of tuberculosis disease phenotype. Thorax 68, 221–229 (2013).

16. Click, E. S., Moonan, P. K., Winston, C. A., Cowan, L. S. & Oeltmann, J. E. Relationship between Mycobacterium tuberculosis phylogenetic lineage and clinical site of tuberculosis. Clin. Infect. Dis. 54, 211–219 (2012).

17. Devaux, I., Kremer, K., Heersma, H. & Van Soolingen, D. Clusters of multidrug-resistant Mycobacterium tuberculosis cases, Europe. Emerg. Infect. Dis. 15, 1052–1060 (2009).

18. Hanekom, M. et al. A recently evolved sublineage of the Mycobacterium tuberculosis Beijing strain family is associated with an increased ability to spread and cause disease. J. Clin. Microbiol. 45, 1483–1490 (2007).

19. Feuerriegel, S., Köser, C. U., Richter, E. & Niemann, S. Mycobacterium canettii is intrinsically resistant to both pyrazinamide and pyrazinoic acid. J. Antimicrob. Chemother. 68, 1439–1440 (2013).

20. Coscolla, M. & Gagneux, S. Consequences of genomic diversity in Mycobacterium tuberculosis. Semin. Immunol. 26, 431–444 (2014).

21. Koya, M. P., Simon, M. A. & Soloway, M. S. Complications of intravesical therapy for urothelial cancer of the bladder. J. Urol. 175, 2004–2010 (2006).

22. Ninane, J., Grymonprez, A., Burtonboy, G., Francois, A. & Cornu, G. Disseminated BCG in HIV infection. Arch. Dis. Child. 63, 1268–1269 (1988).

23. van Soolingen, D. et al. Diagnosis of Mycobacterium microtiInfections among Humans by Using Novel Genetic Markers. J. Clin. Microbiol. 36, 1840–1845 (1998).

24. Kiers, A., Klarenbeek, A., Mendelts, B., Van Soolingen, D. & Koëter, G. Transmission of Mycobacterium pinnipedii to humans in a zoo with marine mammals. Int. J. Tuberc. Lung Dis. 12, 1469–1473 (2008).

25. Brosch, R. et al. A new evolutionary scenario for the Mycobacterium tuberculosis complex. Proc. Natl. Acad. Sci. U. S. A. 99, 3684–3689 (2002).

26. Jagielski, T. et al. Current methods in the molecular typing of Mycobacterium tuberculosis and other mycobacteria. Biomed Res. Int. 2014, 645802 (2014).

27. Steiner, A., Stucki, D., Coscolla, M., Borrell, S. & Gagneux, S. KvarQ: targeted and direct variant calling from fastq reads of bacterial genomes. BMC Genomics 15, 881 (2014).

28. Coll, F. et al. Rapid determination of anti-tuberculosis drug resistance from whole-genome sequences. Genome Med. 7, 51 (2015).

29. Ondov, B. D. et al. Mash: fast genome and metagenome distance estimation using MinHash. Genome Biol. 17, 132 (2016).

30. Bradley, P. et al. Rapid antibiotic-resistance predictions from genome sequence data for Staphylococcus aureus and Mycobacterium tuberculosis. Nat. Commun. 6, 10063 (2015).

31. Nguyen, L.-T., Schmidt, H. A., von Haeseler, A. & Minh, B. Q. IQ-TREE: a fast and effective stochastic algorithm for estimating maximum-likelihood phylogenies. Mol. Biol. Evol. 32, 268–274 (2015).

32. Deatherage, D. E. & Barrick, J. E. Identification of mutations in laboratory-evolved microbes from next-generation sequencing data using breseq. Methods Mol. Biol. 1151, 165–188 (2014).

33. Seemann, T. snippy. (Github, 2015).

34. Maddison, W. P. & Maddison, D. R. Mesquite: a modular system for evolutionary analysis. (2017).

35. Bradley, P. cbg. (Github).

36. Coll, F. et al. A robust SNP barcode for typing Mycobacterium tuberculosis complex strains. Nat. Commun. 5, 4812 (2014).

37. Cingolani, P. et al. A program for annotating and predicting the effects of single nucleotide polymorphisms, SnpEff: SNPs in the genome of Drosophila melanogaster strain w1118; iso-2; iso-3. Fly 6, 80–92 (2012).

38. Stucki, D. et al. Mycobacterium tuberculosis lineage 4 comprises globally distributed and geographically restricted sublineages. Nat. Genet. 48, 1535–1543 (2016).

39. Walker, T. M. et al. Whole-genome sequencing for prediction of Mycobacterium tuberculosis drug susceptibility and resistance: a retrospective cohort study. Lancet Infect. Dis. 15, 1193–1202 (2015).

40. Schleusener, V., Köser, C. U., Beckert, P., Niemann, S. & Feuerriegel, S. Mycobacterium tuberculosis resistance prediction and lineage classification from genome sequencing: comparison of automated analysis tools. Sci. Rep. 7, 46327 (2017).

41. Stucki, D. et al. Two new rapid SNP-typing methods for classifying Mycobacterium tuberculosis complex into the main phylogenetic lineages. PLoS One 7, e41253 (2012).

42. Comas, I., Homolka, S., Niemann, S. & Gagneux, S. Genotyping of genetically monomorphic bacteria: DNA sequencing in Mycobacterium tuberculosis highlights the limitations of current methodologies. PLoS One 4, e7815 (2009).

43. Olea-Popelka, F. et al. Zoonotic tuberculosis in human beings caused by Mycobacterium bovis—a call for action. Lancet Infect. Dis. 17, e21–e25 (2017).

44. Evans, J. T. et al. Cluster of human tuberculosis caused by Mycobacterium bovis: evidence for person-to-person transmission in the UK. Lancet 369, 1270–1276 (2007).

45. Müller, B. et al. Zoonotic Mycobacterium bovis-induced tuberculosis in humans. Emerg. Infect. Dis. 19, 899–908 (2013).

46. Thapa, J. et al. Mycobacterium orygis-Associated Tuberculosis in Free-Ranging Rhinoceros, Nepal, 2015. Emerg. Infect. Dis. 22, 570–572 (2016).

47. Rahim, Z. et al. Tuberculosis Caused by Mycobacterium orygis in Dairy Cattle and Captured Monkeys in Bangladesh: a New Scenario of Tuberculosis in South Asia. Transbound. Emerg. Dis. (2016). doi:10.1111/tbed.12596

48. Dawson, K. L. et al. Transmission of Mycobacterium orygis (M. tuberculosis complex species) from a tuberculosis patient to a dairy cow in New Zealand. J. Clin. Microbiol. 50, 3136–3138 (2012).

49. Kremer, K. et al. Mycobacterium microti: more widespread than previously thought. J. Clin. Microbiol. 36, 2793–2794 (1998).

50. Niemann, S. et al. Two cases of Mycobacterium microti derived tuberculosis in HIV-negative immunocompetent patients. Emerg. Infect. Dis. 6, 539–542 (2000).

51. Xavier Emmanuel, F. et al. Human and animal infections with Mycobacterium microti, Scotland. Emerg. Infect. Dis. 13, 1924–1927 (2007).

52. Horstkotte, M. A. et al. Mycobacterium microti llama-type infection presenting as pulmonary tuberculosis in a human immunodeficiency virus-positive patient. J. Clin. Microbiol. 39, 406–407 (2001).

53. Panteix, G. et al. Pulmonary tuberculosis due to Mycobacterium microti: a study of six recent cases in France. J. Med. Microbiol. 59, 984–989 (2010).

